# OscoNet: Inferring oscillatory gene networks

**DOI:** 10.1101/600049

**Authors:** Alexis Boukouvalas, Luisa Cutillo, Elli Marinopoulou, Nancy Papalopulu, Magnus Rattray

## Abstract

**Motivation:** Oscillatory genes, with periodic expression at the mRNA and/or protein level, have been shown to play a pivotal role in many biological contexts. However, with the exception of the circadian clock and cell cycle, only a few such genes are known. Detecting oscillatory genes from snapshot single-cell experiments is a challenging task due to the lack of time information. Oscope is a recently proposed method to identify co-oscillatory gene pairs using single-cell RNA-seq data. Although promising, the current implementation of Oscope does not provide a principled statistical criterion for selecting oscillatory genes.

**Results:** We improve the optimisation scheme underlying Oscope and provide a well-calibrated non-parametric hypothesis test to select oscillatory genes at a given FDR threshold. We evaluate performance on synthetic data and three real datasets and show that our approach is more sensitive than the original Oscope formulation, discovering larger sets of known oscillators while-avoiding the need for less interpretable thresholds. We also describe how our proposed pseudo-time estimation method is more accurate in recovering the true cell order for each gene cluster while-requiring substantially less computation time than the extended nearest insertion approach.

**Conclusion:** OscoNet is a robust and versatile approach to detect oscillatory gene networks from snapshot single-cell data addressing many of the limitations of the original Oscope method.

## Background

Oscillating genes are expressed in a periodic manner leading to alternating appearance and disappearance of the corresponding mRNA and protein. Oscillatory genes have been shown to play a pivotal role in many developmental processes, by enabling individual systems to implement diverse functions [11]. To identify oscillatory genes a combination of time-lapse microscopy techniques and fluorescent reporter genes is required, which prohibits the assaying of a large number of potential oscillators. In turn, this limits the ability of the experimenter to uncover novel oscillators.

Recent developments in single-cell transcriptomics provide the potential to capture genome-wide transcriptional events. However, single-cell RNA-seq experiments, even when done as time series, pose challenges associated in identifying oscillating genes because cells may be asynchronous in vivo or may lose synchronisation in the time required for sample preparation and sequencing.

To overcome these problems, Leng *et al.* [10] have proposed a computational approach to identify oscillating genes in static, unsynchronised scRNA-seq experiments. They construct a parameterised distance between pairs of putative oscillating genes which is minimised for co-oscillating pairs that have the same temporal profile but different phase. Their approach allows gene-pairs to be ranked by how close they are according to this co-oscillation distance. However, they do not provide a statistical test to decide which gene pairs are oscillatory.

In this contribution, we provide a well-calibrated hypothesis test which allows us to remove the less interpretable cut-off in the first step of the Oscope pipeline [10]. We do this by selecting gene-pairs according to a false discovery rate (FDR) cut-off. Our non-parametric bootstrap approach requires us to optimise the gene-pair distance many times, and therefore we have implemented an efficient optimisation algorithm to minimise the gene-pair distance function. We then construct an undirected network of co-oscillating genes based on all gene-pairs which pass the FDR threshold. Community extraction and network analysis methods can then be used to identify statistically significant co-oscillating groups of genes. We also discuss three distinct statistical tests that can be employed to assess the significance of different aspects of the inferred network communities, namely community distinctiveness and enrichment, in terms of both genes (graph vertices) and their co-oscillations (graph edges). We conclude with a description of a simple method to infer pseudo-time for each gene community, inferring a cyclical ordering on a low-dimensional projection.

## Review of the Oscope method

The Oscope method [10] uses a paired-sine model and K-medoids clustering to identify groups of oscillatory genes. For each oscillatory group, an extended nearest insertion algorithm is used to construct the cyclic order of cells, defined as the order that specifies each cell’s position within one cycle of the oscillation of that group [1]. The method requires normalised gene expression and the use of only high mean, high variance genes is recommended [1].

The method relies on the computation of a pairwise gene expression similarity. Let two genes be *X* = sin (*ωt* + *φ*) and *Y* = sin (*ωt* + *φ* + Ψ_*XY*_), that is they have identical profiles except for a phase shift Ψ_*XY*_. For *N* cells the co-oscillation distance is *d* (*X, Y |*Ψ_*XY*_):

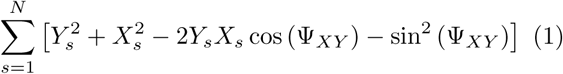

The derivation is provided in the supplementary material (Section 3). The assumed sinusoidal form is not a major restriction, given that the time *t* is not required in the co-oscillation distance^[1]^. This is the crucial step in the algorithm as it allows the gene-pair similarity to be estimated without knowledge of the pseudo-time.

A limitation of this approach is that the cell population is assumed to be homogeneous. Including cell populations that are not relevant to the process under investigation will result in reduced accuracy as the co-oscillation distance will be averaged across both informative and non-informative cells. We therefore recommend irrelevant cell populations are removed prior to the application of the Oscope and OscoNet methods. This can be accomplished by visualisation and clustering methods such as t-SNE and k-means to identify cell populations and using gene markers or other information to characterise each cell cluster.

In the original Oscope approach, only genes that appear in the top *T* % of the ranked pairwise similarity list are selected for the subsequent K-medoids clustering step. The choice of *T* can have a significant impact on downstream analysis. However, the optimal choice of *T* is not obvious and will likely differ depending on characteristics of the data, e.g. number of co-oscillatory pairs, technical/biological noise levels, number of cells in the dataset etc. In our work, we instead propose to use a well-calibrated non-parametric test to select gene-pairs with a given false discovery rate (FDR). Genes that contribute to these gene-pairs are then selected for further downstream analysis.

A further limitation of the Oscope approach is that genes which are linearly correlated cannot be disambiguated from genes pairs with 0 or *π* phase shift. Therefore a heuristic is used to remove clusters that have a high proportion of similarities with 0 or *π* phase shift.

Finally, the extended nearest insertion (ENI) algorithm is used to estimate pseudo-time per gene cluster. The use of the ENI algorithm is not critical to the algorithm and can be replaced by a computationally more efficient probabilistic approach as we describe below.

## Methods

Our overall approach is summarised in Figure 1. Following a quality control step (see Appendix), we propose three main phases, namely: building a network of gene co-oscillations using a non-parametric hypothesis test, identify the statistically significant communities in the resulting network and infer pseudo-time for each gene community.

**Figure 1.**
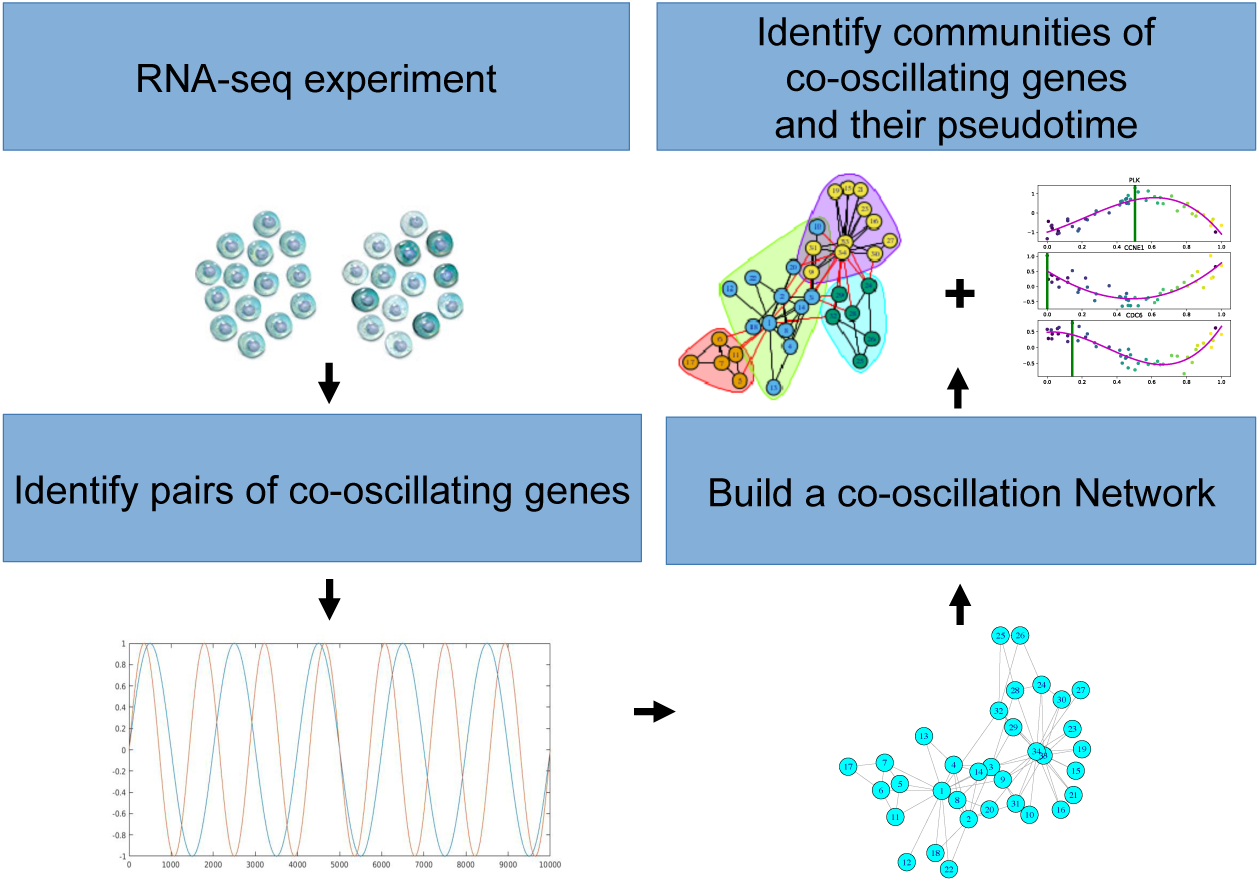
Overall approach: A statistical test is used to identify co-oscillatory gene-pairs at a given FDR threshold. Genes contributing to these gene-pairs are included in a co-oscillation network. The q-value from our co-oscillation test provides an estimate of the number of false links in this network. We can then identify communities in this network and infer the pseudo-time assciated with each community.

### Non-parametric test on co-oscillation

The key insight in the Oscope algorithm is that a gene-pair oscillation score can be defined without requiring a prior estimation of pseudo-time. Rather than specifying a threshold on the number of genes to keep, here we develop a non-parametric hypothesis test to assess the significance of two genes co-oscillating.

We employ a bootstrap approach which randomly permutes the order of cells for one of the genes in the gene-pair distance estimation in order to perform a statistical test. The cell index is permuted for one gene in a pair and we calculate the resulting pairwise gene similarity *d* (*X*, 𝒫(*Y*) Ψ_*XY*_) (recall Equation (1)) where 𝒫(*Y*) is the permuted cell order for gene *Y*. For each randomisation, we estimate the phase shift by minimising the pairwise distance,

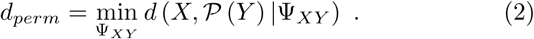

The p-value estimate for the gene pair *XY* is the fraction of times *d_perm_ < d*. Specifically for *B* randomisations the bootstrap p-value is

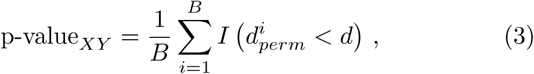

where 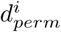 the *i*^th^ randomisation, *d* = *d* (*X, Y |*Ψ_*XY*_) the unpermuted pairwise distance and *I*(⋅) the indicator function, which is 1 when the inequality is true and 0 otherwise.

In our approach we need to correct for the effect of multiple hypothesis tests since we are testing many gene-pairs. We have selected to use the q-value approach of [18] which controls for the false discovery rate (FDR). The simple Bonferroni correction is too conservative in the case of very large numbers of tests and is also based on an unrealistic assumption that tests are independent. The q-value is an adjustment of the traditional p-value where the FDR is minimised based on the distribution of the p-values across all the tests performed. The q-value is an estimate of the FDR at a given p-value threshold obtained from modelling background (null) and signal contributions to the p-value distribution. This approach does not assume independent tests and has been shown to perform well in applications with very large numbers of tests. The q-value provides an estimate of the number of false edges in our co-oscillation network.

As the bootstrap approach is computationally demanding, we have expressed the distance measure using matrix operations and we provide an efficient implementation. We have implemented the bootstrap test in Python using NumPy vector operations and we have also implemented a faster version in Cython, which allows C code to be called from Python. A performance comparison of the different implementations is given in Figure 2 wherein we compare our implementation of the pairwise distance calculation (NumPy, Cython) to the standard Oscope [10] R implementation. As all three implementations are able to leverage multi-core parallelism, our test is performed across 12 CPU cores. We see that both Numpy and Cython are substantially faster than the iterative R version which uses the Oscope bioconductor package. The Numpy and Cython implementations achieve an average speed-up of 6.5 and 93 respectively. The significant performance improvement allows for the practical use of the bootstrap algorithm. The computational penalty for a fully nonparametric significance test is therefore minimised.

**Figure 2.**
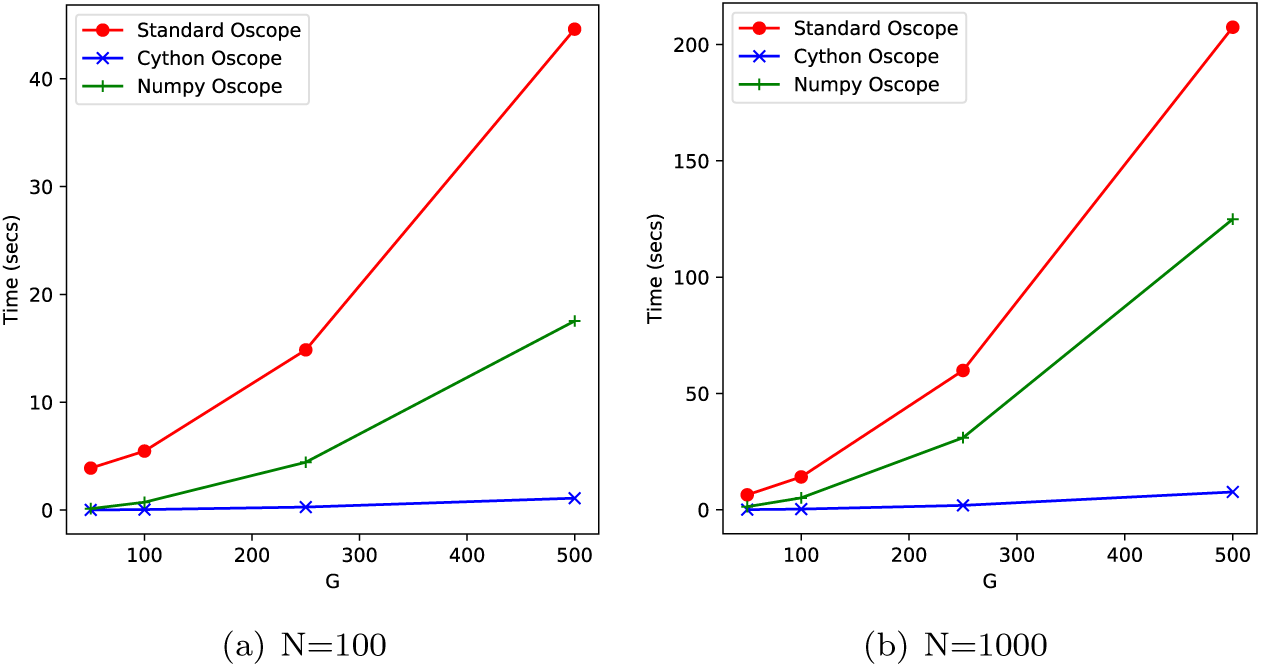
Performance comparison of different Oscope implementations on computing the pairwise oscillating distance. Vertical axis is elapsed time in seconds and horizontal axis is number of genes (G). Standard Oscope Leng *et al*. [10] is an R implementation. All methods use 12 cores and the minimum elapsed time from 2 runs is reported.

### Network analysis

We use the gene-pair test to construct a network of genes in which connections denote statistically significant co-oscillation. In the network setting, a variety of algorithms have been developed to extract communities within the entire network. The problem of community detection, or graph partitioning, in a network has been widely studied in a variety of research fields. In the present work we consider community extraction algorithms based on maximising modularity, a measure of community distinctiveness. The modularity, often referred to as *Q*, was defined by Newman and Girvan [13] as the fraction of the edges that fall within a given groups of nodes (community partition) minus the expected fraction if edges were distributed at random. The concept of modularity is based on the idea that a random graph is not expected to have a cluster structure. In the *Synthetic study* section, we compare three popular community extraction algorithms^[2]^, namely: walktrap [14], fast-greedy [3] and infomap [16], and find the walktrap algorithm to be the most robust.

Once our network is estimated, it is still a complex network and we need to take a sense out of it. A common approach to address this issue is finding an underlying community structure, if it exists [6]. The identified Communities will allow us to create a large scale map of our network [4]. We aim to use individual communities to shed light on the function of the co-oscillation represented by the network itself. Indeed, in biological networks, communities often correspond to functional units of a system. For example in metabolic networks [9], such functional groups correspond to cycles or pathways whereas in the protein-protein interaction networks [7], communities correspond to proteins with similar functionality inside a biological cell.

To assess significance of each community, we employ three statistical tests that examine different aspects of the community structure. Firstly we apply a Wilcoxon test to compare the distribution of edges inside each community to the distribution of edges that have crossed the community boundary, i.e. one node connected within the community and the other without. This tests allows to assess how well-defined a community is and complements the maximisation of the modularity objective during the community extraction stage.

We then validate our results cross-referencing with a list of known oscillating targets. To this end, for each identified community, we perform a set of gene enrichment tests with respect to Cell Cycle (CC) related genes (Gene Ontology term GO:0007049). Firstly, we test each community for standard gene enrichment using a hypergeometric test; this test checks whether the genes in every single community are significantly enriched in the list of CC related genes. However this test only considers membership to a community and does not take into account the within community connections corresponding to gene co-oscillations. For a more stringent test we also employ a network enrichment test [17]. For every community that contains CC related genes, we are performing a network enrichment test between two lists of genes, namely: list *A* - the list of CC related genes in the community, list *B* - the list of all the genes in the community. This test aims at looking at the connection between our target genes to see if their links are statistically significant, hence if there is an enrichment. The main idea behind this test is that, under the null hypothesis of no enrichment, the number of links between two gene sets A and B follows an hypergeometric distribution. This test considers the connectedness among CC related genes within a single community. Therefore it may be more stringent than the hypergeometric test, given that CC enriched communities where the related CC genes are only connected through intermediaries are not found to be directly co-oscillating. In our results, we show the validation of communities in terms of the Wilcoxon test and the CC enrichment in terms of network enrichment test [17].

We also introduce an index, community density, expressing the fraction of the actual number of edges in a community with respect to the total number of edges in the network. We also provide a relative community density, defined as the fraction of edges in a community given the maximum number of edges feasible in a community of that size.

### Pseudo-time estimation

We use Laplace Eigenmaps [2] to reduce the gene expression space to a two dimensional latent representation. Laplace Eigenmaps, also known as spectral embedding, is a non-linear dimension reduction method using a spectral decomposition of the graph Laplacian to preserve local distances in the gene expression space. The method is fast to compute as it involves a partial eigenvalue decomposition of a sparse graph matrix, namely the graph Laplacian. Further, the minimisation problem has a unique solution and therefore does not suffer from local minima. As Belkin and Niyogi [2] discuss, the method is robust to outliers and noise as only local distances are used to estimate the neighbourhood graph.

The method consists of

1. Graph Construction: create sparse adjacency matrix *A* using *n* nearest neighbours.
2. Graph Laplacian Construction: *L* = *D − A*, where *D* is the degree matrix, the number of edges attached to each vertex.
3. Partial Eigenvalue Decomposition. Find top-*Q* eigenvalues of sparse matrix where *Q* is the dimensionality of the latent space. In our case this is always *Q* = 2.

The critical parameter in the algorithm is *n*, the number of nearest neighbours. Our pseudo-time estimation algorithm consists of a search on a grid of values specified by a minimum *m* and maximum value *M*. For *n ∈ {m,…, M*}:

1. Compute Laplacian eigenmap on 2 dimensions.
2. pseudo-time: compute angle in radians [−*π, π*] by arc tangent. Convert to [0, 1].

The latter step imposes a periodic constraint on the pseudo-time: pseudo-time is defined as the angle of a cell in a circle.

To select the best value for the number of neighbours *n*, we apply a probabilistic periodic model to assess each pseudo-time for all genes in the cluster. We employ a Bayesian Gaussian process latent variable model [20] with a one-dimensional periodic kernel. All hyperparameters are optimised except the pseudo-time. We use 25 inducing points to ensure efficient optimisation and calculation of the model likelihood. We then use the marginal model likelihood to select the best pseudo-time and the number of neighbours *n*.

## Results

In order to validate our overall approach and compare to the original results in [10], we evaluate the ability of OscoNet to identify oscillating groups of genes by applying it both to synthetic data, a microarray time-series study of oscillating genes [21] and to the profiles of single undifferentiated human embryonic stem cells (hESCs) [19]. For the biological data, we evaluate the enrichment of the discovered clusters with respect to the Gene Ontology Cell Cycle (CC) biological process related genes (Gene Ontology term: 0007049).

### Synthetic study

We generate a synthetic dataset of 1,000 genes and 100 cells following the setup in Leng *et al.* [10]. We simulate 1000 genes of which 180 are samples from a sinusoidal function with additive Gaussian noise. The 180 oscillators are simulated in 3 frequency groups, each group containing 60 genes. The relative frequencies of the 3 groups are proportional to 2:3:6.

For each group, genes were further simulated having strong and weak signals. Half of the oscillatory genes were simulated as strong oscillators with noise variance *σ*^2^. The other half were simulated as weak oscillators with 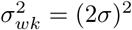. The starting phase Φ varies in different genes within a frequency group. The remaining genes were simulated as independent Gaussian noise.

### Hypothesis test

We calculate the false discovery rate on all gene pairs, defined as:

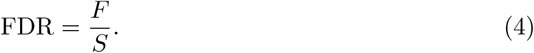

where *S* is the estimated number of significant gene pairs and *F* the number of false positive pairs. This is a more stringent criterion than in [10] where the number of oscillating genes was used and not co-oscillating pairs.

The accuracy of the non-parametric test with different number of bootstrap samples and the original Oscope method is shown in Figure 3 (a) for noise level of *σ*^2^ = 0.05. When considering higher noise levels the results are similar (see supplementary). When a very small number of bootstrap samples is used (*B* = 100) the test is not well-calibrated. However when the number of samples is increased the bootstrap test better approximates the true FDR.

**Figure 3.**
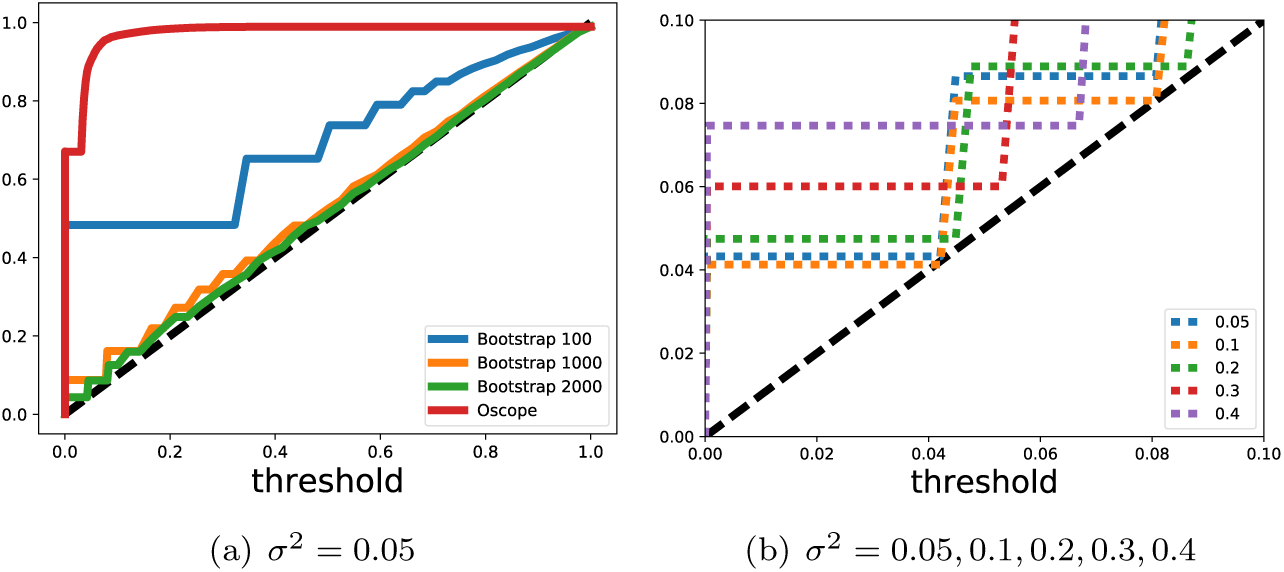
False discovery rate (FDR) for the synthetic study. The synthetic data was generated using noise variances σ^2^ = [0:05, 0:1, 0:2, 0:3, 0:4]. Vertical axis is achieved FDR and horizontal axis the significance level used. Plot (a) gives the FDR for noise level σ^2^ = 0.05 for different number of bootstrap samples and the Oscope method for different cut-offs. Plot (b) compares the effect of different noise levels on the FDR of the bootstrap hypothesis test using 2000 samples. Magnus: put achieved FDR on y-axes and q-value on x-axese here

However in all cases, we observe an elbow effect when decreasing the desired level of FDR. At a certain point of significance, the observed FDR is not further reduced as we decrease the desired FDR. This occurs at increasingly smaller thresholds as the number of bootstrap samples is increased. However we suspect that the number of cells and the variability of gene expression also play a role; when increasing the number of cells for instance, statistical power is expected to increase as we get higher resolution on the tails of the null distribution. Another factor is the amount of noise in the data as is demonstrated in Figure 3 (b) where we show the effect of different noise levels on the threshold value of the elbow after which no improvement in FDR is observed. As the noise level decreases, the statistical test is able to achieve lower FDR.

A possible improvement on our test would be to introduce an explicit model for the tails of the null distribution. Such a semi-parametric bootstrap approach would allow for lower achievable FDR levels as long as the modelling assumptions are accurate.

The poor performance of standard Oscope can be understood by the nature of the threshold used. At the lowest noise level (0.05) the method correctly identifies all the co-oscillating gene pairs at the default 5% threshold achieving a perfect true positive rate (*TPR* = 1.0) as does the OscoNet hypothesis test. However the former gets a false positive rate of 10% (*FPR* = 0.10) while the latter achieves a lower rate of 0.1% (*FPR* = 0.001). The resulting FDR for Oscope is 0.91 and for OscoNet is 0.09 reflecting the highest sensitivity of the latter. The large FPR achieved by Oscope demonstrates the deficiency of setting such a threshold. In the supplementary (Section 1), we present the FDR, FPR and TPR achieved across all noise levels.

Note that a bootstrap test is implemented in the Oscope pipeline after the clustering step to flag clusters that should be removed prior to downstream analysis. As this test is applied after the threshold of the top ranked pairs of genes, the choice of threshold has significant impact on the results. In addition, by applying the bootstrap test on the cluster level, an entire cluster is either kept or removed whereas our approach maintains all significant gene pairs. We argue our approach is logically more coherent, since the clustering should be applied on only significant gene pairs rather than testing for significance after the clustering is completed.

### Comparing community extraction methods

In order to validate our approach, we want to find densely connected subgraphs, also called communities, in the constructed synthetic graphs. To this aim we compare three well known network clustering algorithms available in the *R* package *igraph*, namely: *walktrap*, *fastgreedy* and *infomap*. In *walktrap* the communities are assigned via random walks [14]. The idea is that short random walks tend to stay in the same community. In *fastgreedy*, the communities are estimated via directly optimising a modularity score [3]. On the other end, *infomap* aims to find a community structure that minimises the expected description length of a random walker trajectory [16]. We refer to the original papers for a detailed description. In an ideal case, true simulated oscillating genes should be clustered in the same community. For each of the five noise levels, as defined in the *Synthetic study* section, and for each one of the three community detection algorithms, we compare the recovered network partition to the true underlying one computing the *Adjusted Rand index* (*ARI*). This measure is a scalar comparing two partitions on the same network. The *ARI* index, first introduced in Hubert and Arabie [8], has zero expected value in the case of random partition, and it is 1 in the case of perfect agreement between two partitions. As depicted in Figure 4, the index decreases when the level of noise increases and the most stable results are achieved by the *walktrap* algorithm. We therefore select the *walktrap* as the community extraction method in the clustering step of our overall approach.

**Figure 4.**
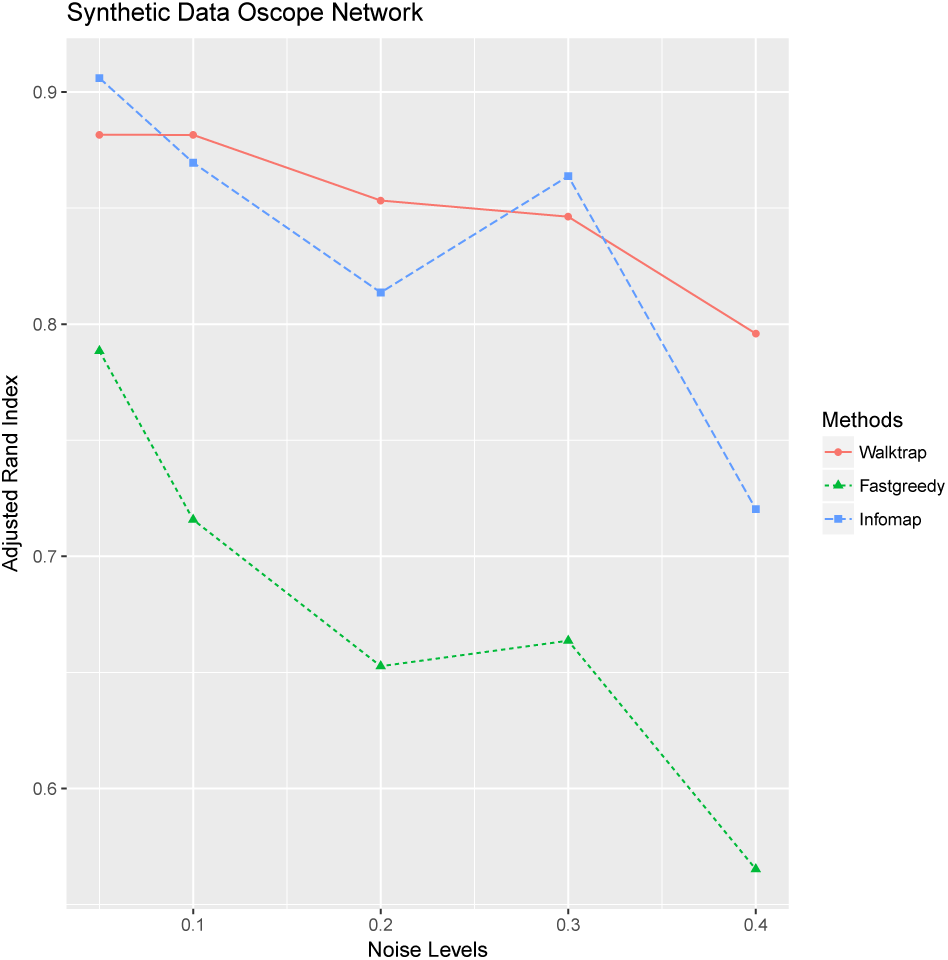
Comparison of three clustering algorithms on the synthetic network. For each method the Adjusted Random Index (vertical axis) is provided at different noise levels (horizontal axis).

### cDNA microarray time course

We applied OscoNet to a microarray time-series study of oscillating genes studied in [21].

Following Leng *et al.* [10], we use experiment 3 from the original study [21] where double thymidine block was used to synchronise HeLa cells which were profiled at 48 time points following synchronisation. We start with a set of 41508 transcripts containing 1134 cell cycle (CC) oscillators [21]. This corresponds to a set of 9559 genes containing 874 oscillating genes [21].

We filter transcripts using variance based filtering with a cut-off of 0.9 [5] This results to 4151 probes containing 360 CC oscillators. This filtering step is more stringent than the default test on mean and variance of expression used in the Oscope paper. This filtering step also ensures that the set of transcripts we consider does not include high error measurements and includes the most informative genes only. In both cases, the data is normalised to [−1, 1].

Finally, we identify a total of 3884 genes that have at least one significant co-oscillation where the total number co-oscillations is 185,548. This corresponds to a highly sparse network; specifically the sparseness is 0.0123 calculated as the ratio of the number of extant network edges to the maximum number possible. All the 60 CC oscillators are present in the network.

In the following, we summarise the results when applying the standard Oscope approach or respectively our approach. The Oscope approach was applied to the exact same data using the same normalisation. We apply the standard paired sine model and only gene pairs with similarity score in the top 5%. This corresponds to the default setting recommended in Oscope. We apply K-medoids with the maximum number of clusters set to *K* = 5. Finally, we remove clusters flagged by Oscope as having mostly linear effects.

The application of standard Oscope led to the identification of two significant clusters of transcripts: one of size 72, of which 70 are cell cycle related and one of size 136 respectively, of which 13 are cell cycle related. We note that no clusters were removed by the linear filtering step of the Oscope pipeline.

The application of OscoNet to the same dataset led to the identification of 5 significant clusters, according to the Wilcoxon test (significance *α* = 0.01) described in the previous section. Singleton communities with a single gene are excluded.

In Table 1 we show all the significant clusters according to the Wilcoxon test ranked by the relative density. Of these 5, 3 have at least one CC transcript. Of those 3, only 1 is enriched according to the NEAT and hypergeometric tests with *α* = 0.01.

**Table 1.** Microarray data: Relative ratio refers to the percentage of CC transcripts in the community.

| Size | CC | Density | Relative Density | Neat | hypergeometric |
| --- | --- | --- | --- | --- | --- |
| 27 | 0 | 0.00 | 0.44 | N/A | N/A |
| 265 | 239 | 0.08 | 0.41 | $< 1e - 6$ | $< 1e - 6$ |
| 1157 | 99 | 0.80 | 0.22 | 0.28 | 0.83 |
| 131 | 0 | 0.01 | 0.17 | N/A | N/A |
| 82 | 1 | 0.00 | 0.11 | 1.00 | 1.00 |

The only CC enriched community has 239 CC genes out of 265 and is the second-ranked community as per relative density. This community is well-separated from the network and is enriched in CC from both enrichment perspectives, namely traditional enrichment in terms of genes and the network interconnections between the CC genes present in the community.

Importantly, this community includes the 70 CC related transcripts identified as co-oscillating by Oscope. Therefore this community is a more detailed and richer description of the CC processes than that uncovered by Oscope.

#### Pseudo-time

We evaluate the performance of the extended nearest insertion (ENI) method of [10] and spectral methods on estimating pseudo-time. We run the ENI method using 30000 iterations, with a sliding polynomial of degree 3 and 4 starting points for the polynomial fitting. The pseudo-time order is not changed after approximately 6500 iterations in our experiments even when setting the maximum number of iterations to 100k. The ENI algorithm took on average 5-7 hours in our experiments while the spectral method takes less than 1 minute including the time to search over the parameter space. This makes the method very easy to use in practice in addition to the superior performance which we now discuss.

We assess the performance of the method when trained on the 72 gene cluster defined by Oscope and the 265 gene cluster defined by OscoNet. We also evaluate the performance of the pseudo-time on both of these clusters.

The dimensionality reduction and circle estimate by the spectral method for the two gene clusters is shown in Figure 5. In both cases the cells can be well described by a circular path in the 2-D latent space projection of the data.

**Figure 5.**
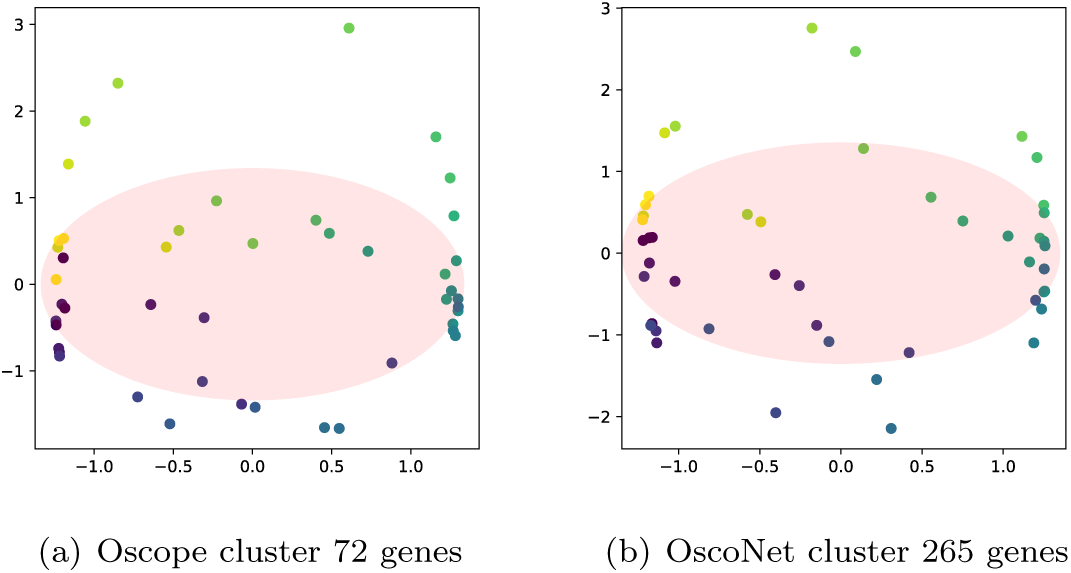
Microarray data: spectral dimensionality reduction trained on the two different clusters. Each cell-dot is coloured by the true base cycle stage. Axis denote the two latent dimensions estimated using the spectral method.

Our main validation measure is the Spearman correlation of the estimated peak times in the reconstructed gene expression to the ground truth (Table 2). The ground truth has been constructed by folding the time series in Whitfield *et al.* [21] along every period. ENI provides consistently poor results despite the large iteration count (30k iterations). The spectral method is simple and provides good results. It can also be easily extended in an intuitive manner to cover more complex cases - see the Discussion section. Comparing the pseudo-time achieved by the different clusters using the spectral method, we note that using the larger gene cluster identified by OscoNet (265 genes) results in a more accurate pseudo-time. This is reflected in the higher peak time rank correlation; the larger cluster estimated by OscoNet is more informative on the true time compared to the smaller cluster estimated by Oscope. In the supplementary (Section 2), the estimated and actual peak times for each gene are shown.

**Table 2.** Microarray data: spearman rank correlation of peak times.

| Method | Training | Predicting cluster peak time |  |
| --- | --- | --- | --- |
|  |  | 72 | 265 |
| Spectral | 72 | 0.94 | 0.81 |
|  | 265 | 0.99 | 0.83 |
| ENI | 72 | 0.17 | 0.28 |
|  | 265 | -0.19 | 0.14 |

We also report on the roughness measure of the true time series and the reconstructed version. This encapsulates how smooth a time series is by comparing neighbouring values with smaller values corresponding to a smoother time series; see Appendix for details. In Table 3 we report the difference of the median roughness of the estimated pseudo-time and true time. Overall the ENI results in an increase in the roughness measure while the Spectral method results in lower values reflecting smoother time courses. The spectral method achieves similar performance under both the smaller Oscope cluster (72 genes) and the larger OscoNet cluster (265). Therefore the differences reported above in peak time accuracy are not caused by smoother time courses.

**Table 3.** Microarray data: change in median roughness.

| Method | Training | Predicting gene set |  |
| --- | --- | --- | --- |
|  |  | 72 | 265 |
| Spectral | 72 | -0.20 | -0.09 |
|  | 265 | -0.19 | -0.10 |
| ENI | 72 | 0.50 | 0.38 |
|  | 265 | 0.56 | 0.39 |

In Figure 6 we show the gene expression for three genes using the experimental time. In Figures 7 we show the reconstructed base cycle and estimated gene expression. The pseudo-time methods have been estimated using the larger OscoNet cluster. As expected, the ENI profiles are poor due to the poor pseudo-time estimate. The spectral pseudo-time is anticorrelated and therefore provides a mirror image of the true time series profile.

**Figure 6.**
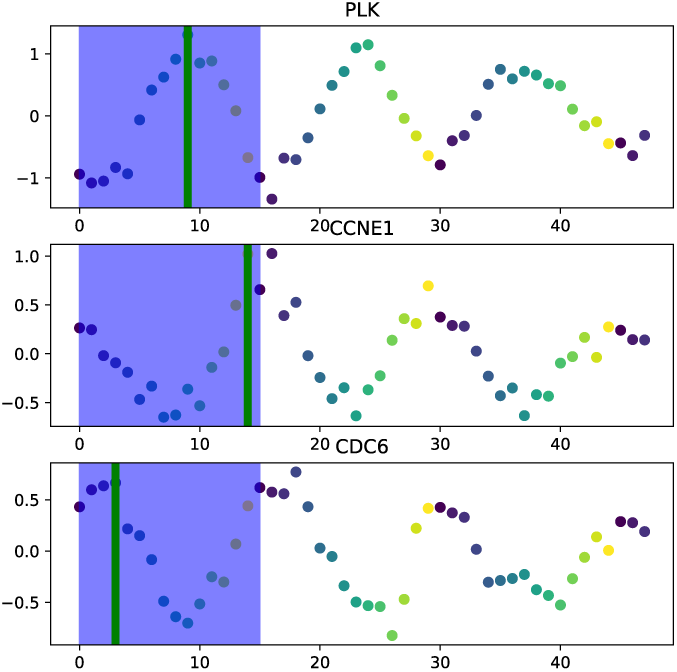
Microarray data: oscillatory gene expression for CDC6, PLK and CCNE1 genes. The true cyclical expression is shown. The length of the base cycle is highlighted in blue background.

**Figure 7.**
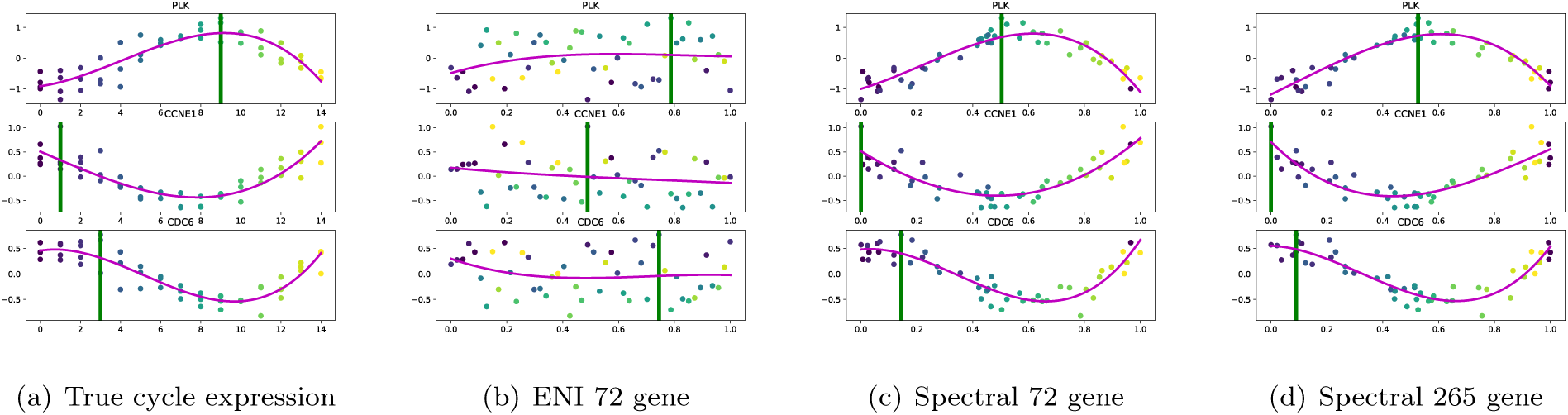
Microarray data: oscillatory gene expression for CDC6, PLK and CCNE1 genes. The cyclical expression and the Oscope-ENI and spectral approach reconstructions are shown. The green vertical line denotes the peak oscillation time.

### Single-cell embryonic stem cells

To evaluate OscoNet on scRNA-seq data, we analysed profiles of single undifferentiated human embryonic stem cells (hESCs) as described in [19]. In particular, we analysed three replicate scRNA-seq experiments on H1 hESCs, with a sample size *n* = 213.

We apply our method to *n* = 213 samples with *G* = 2375 genes following the same steps as [10] in terms of quality control. In particular, genes have been filtered by mean level and variance as in Leng *et al.* [10]. A total of 1914 genes have at least one significant co-oscillation with 5057 co-oscillatory pairs found. The sparseness value for the network is 0.001, an order of magnitude smaller than the previous study suggesting a sparser network.

A total of 677 genes are used to test CC enrichment (term GO: 0007049). A total of 153 CC genes are present after the quality control step, 134 of which are subsequently present in the co-oscillatory network.

The original algorithm in Oscope identified 29 genes as co-oscillating, 21 of which are annotated as belonging to the Gene Ontology Cell Cycle biological process. The Oscope algorithm was run with default values as reported in Leng *et al.* [10]. In total the k-medoids Oscope step finds 5 clusters, 2 of which are eliminated by the subsequent linear filtering step. The CC enrichment of each cluster is given in Table 4.

**Table 4.**
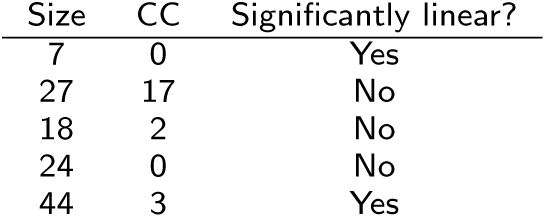
Single-Cell RNA-seq: CC enrichment for each Oscope cluster.

| Size | CC | Significantly linear? |
| --- | --- | --- |
| 7 | 0 | Yes |
| 27 | 17 | No |
| 18 | 2 | No |
| 24 | 0 | No |
| 44 | 3 | Yes |

After applying OscoNet, we assess the significance of each community using a Wilcoxon test as described previously. The results are shown in Table 5. Hence we identify 6 significant communities using the Wilcoxon test (*α* = 0.01). As in the previous study, we eliminate singleton communities that contain only one gene.

**Table 5.** Single-Cell RNA-seq: Relative ratio refers to the percentage of CC transcripts in the community.

| Size | CC | Density | Relative Density | Hypergeometric | Neat |
| --- | --- | --- | --- | --- | --- |
| 4 | 1 | $< 1e-4$ | 0.50 | 0.01 | 1.00 |
| 6 | 0 | $< 1e-4$ | 0.33 | N/A | N/A |
| 77 | 32 | 0.09 | 0.15 | $< 1e-6$ | $< 1e-6$ |
| 155 | 7 | 0.14 | 0.06 | $< 1e-6$ | 0.48 |
| 188 | 11 | 0.10 | 0.03 | $< 1e-6$ | 0.16 |
| 234 | 17 | 0.15 | 0.03 | $< 1e-6$ | 0.03 |

Only one community is both node and edge enriched according to the hypergeometric and NEAT tests (*α* = 0.01). This community is enriched in CC with 32 genes out of 77 belonging to the term GO: 0007049. This community contains the standard Oscope cluster of 29 genes reported in Leng *et al.* [10]. As in the previous study, we find the OscoNet approach is more sensitive than Oscope identifying a cluster with a superset of CC genes.

The two highest ranked communities in terms of relative density have a small size of only 4 and 6 genes respectively. These are ranked highly because of their exceptionally small size and are not CC enriched. A threshold on the community size would exclude such small communities.

#### pseudo-time

We compare pseudo-time using the 77 and 29 gene clusters estimated by OscoNet and Oscope respectively. ENI takes an average of 13 hours of computing time to estimate the pseudo-time for both clusters while the spectral method requires approximately 30 minutes. The difference in computational time is that ENI is leveraged the 2-opt algorithm to reduce the likelihood of local minima. In contrast, the spectral method has a unique solution for a given number of neighbours *n* and is quick to evaluate leveraging efficient sparse eigenvalue solvers. The grid search we perform over different values of *n* ensures the robustness of the results.

In Figure 8 we show the fit for the spectral method for both gene sets. There is more noise in the data than in the previous case study, and the pattern is less clearly circular. Despite this, the algorithm is able to recover the cell cycle order as we now show.

**Figure 8.**
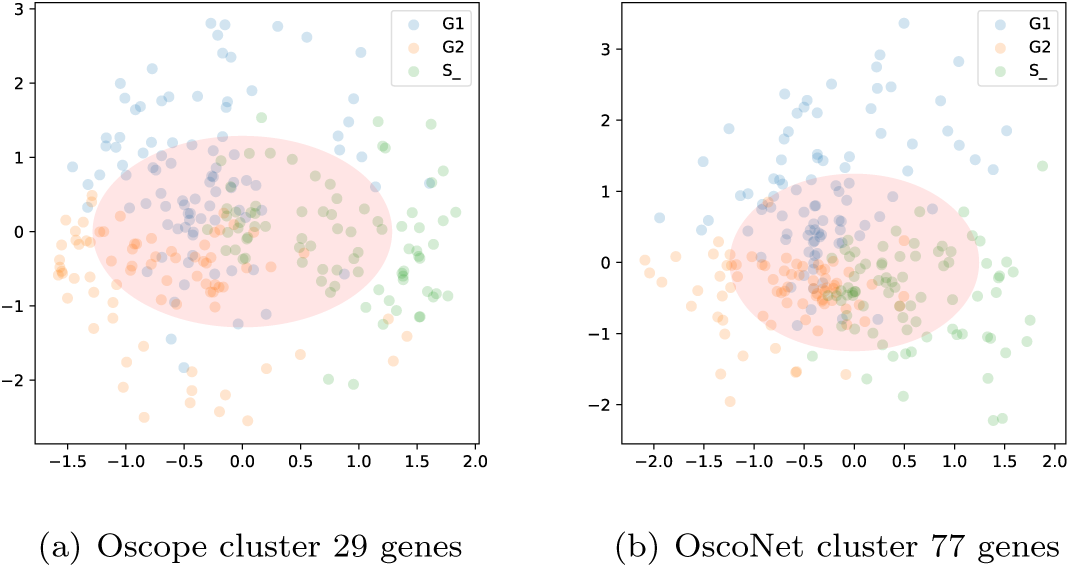
Single-cell RNA-seq: spectral dimensionality reduction trained on the two different clusters. Each cell-dot is coloured by the true base cycle stage, namely G1, G2 and S phase.

We compare the performance of the algorithm estimating pseudo–time using all 460 cells and assessing the separation of cell cycle stages on the labelled subset of cells which consists of 247 cells. After pseudo– time estimation we apply the k-means algorithm with *K* = 3 to identify the three clusters associated with G1, G2 and S phase. We compute the adjusted rand index to contrast the clustering results with the cell cycle labels; the ARI measure is the range [0, 1] with higher values reflecting better agreement between the estimated and true cell cycle stages.

In Table 6 we show the ARI results to the spectral and ENI method. The performance of the latter is poor which can be validated when looking at the gene expression output (Figure 9). The spectral method provides better accuracy for both gene clusters with an ARI of 0.43 when using the Oscope 29 gene cluster and 0.49 when using the OscoNet 77 gene cluster. The higher ARI for the latter, reflects the richer information content of the OscoNet cluster compared to the Oscope cluster; the OscoNet spectral ordering contains fewer cells assigned to an incorrect cell cycle stage than the Oscope spectral ordering as is evident in Figure 9 for four example genes.

**Table 6.** Single-cell RNA-seq: Adjusted Rand Index (ARI) on *K* = 3 K-mean clustering vs true cell-cycle stage.

| Method | Training | ARI |
| --- | --- | --- |
| Spectral | 29 | 0.43 |
|  | 77 | 0.49 |
| ENI | 29 | 0.05 |
|  | 77 | 0.02 |

**Figure 9.**
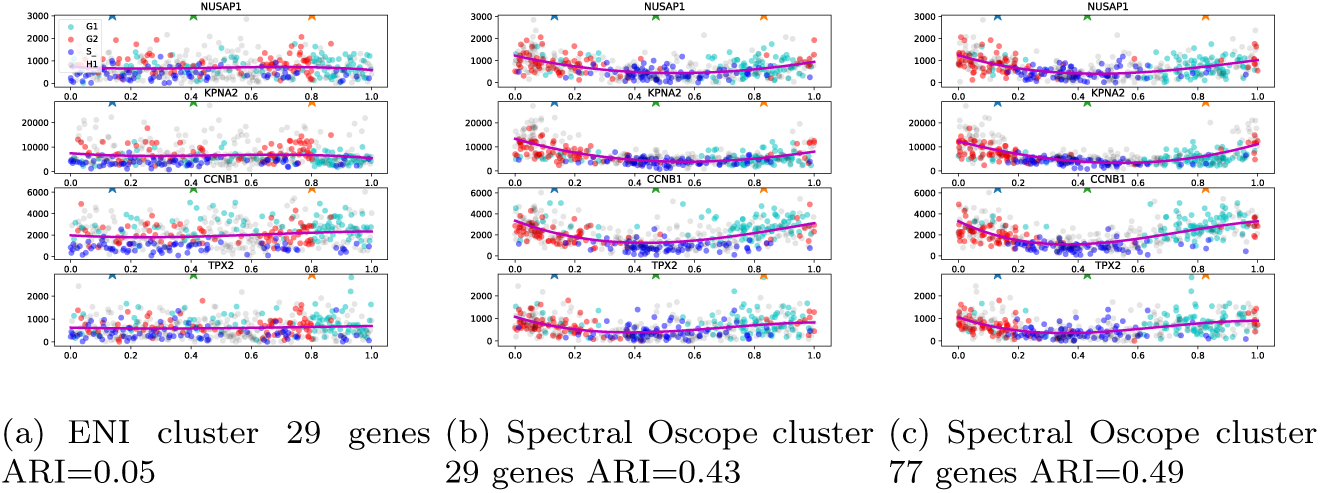
Single-cell RNA-seq: Comparing pseudo-time estimation using ENI and the spectral method. Cells have been coloured by the cell cycle stage or as grey when not known.

The poor performance of the ENI algorithm is confirmed when looking at the roughness value (Table 7). The ENI algorithm has higher roughness values for all combinations examined while the spectral method achieves similar roughness values with both clusters.

**Table 7.** Microarray data: change in median roughness.

| Method | Training | Predicting gene set |  |
| --- | --- | --- | --- |
|  |  | 29 | 77 |
| Spectral | 29 | 1.25 | 1.35 |
|  | 77 | 1.23 | 1.35 |
| ENI | 29 | 1.41 | 1.41 |
|  | 77 | 1.42 | 1.40 |

## Discussion

We have presented a well-calibrated hypothesis test that avoids a need for a threshold cut-off on the pairwise distance matrix. Further, our implementation is highly scalable as it employs a vectorised parallel implementation that was demonstrated to be orders of magnitude faster than the current Oscope implementation.

We have then performed network analysis to identify statistically significant communities of genes again avoiding any need to set the maximum number of communities as is needed in the K-medoids approach of Leng *et al.* [10]. We have also been able to select the significant communities based on a standard statistical test and assess the significance of enrichment in terms of both genes and their co-oscillatory relationship. We have validated our approach on synthetic data demonstrating that it is well-calibrated with respect to the false discovery rate unlike Leng *et al.* [10].

We contrasted our approach to that of [10] on both microarray and single-cell RNA seq data with known oscillators. In both cases, we are able to recover the original Oscope cluster of genes in addition to discovering more oscillators.

We estimate a unique pseudo-time per gene community using a spectral approach that is is straightforward to apply, requires an order of magnitude less computation than the ENI approach proposed in Leng *et al.* [10] and is more accurate as reflected in both peak time correlation in the microarray data and cell cycle stage separation in the single-cell data.

One issue with our current approach is that for low number of cells, the small number of possible permutations in the similarity measure reduces the power of the bootstrap hypothesis test resulting in a high minimum achievable FDR. A semi-parametric bootstrap approach should help increase the power of the test for small number of cells and low FDR that is typically required.

Finally, extending this approach as a probabilistic model including effect of dropout which is prominent in single-cell datasets, may further increase the accuracy of the method and allow for quantification of uncertainty in the resulting co-oscillating gene clusters.

The pseudo-time model could be extended to use a probabilistic model to jointly estimate pseudo-time while-imposing a periodicity constraint. This will also allow for selecting the number of latent dimensions by maximising the model evidence. A probabilistic formulation of Laplace Eigenmaps has already been described in Lu *et al.* [12] where the Laplacian Eigenmaps latent variable model is described. In future work, one could extend this class of model, in the probabilistic or deterministic formulation, to include a periodic constraint.

The method we have presented can be incorporated in a single-cell analysis pipeline to allow biologists to uncover novel periodic dynamic gene regulatory mechanisms. Our approach avoids less interpretable cut-offs and relies on well-calibrated statistical tests. As the availability and size of single-cell whole-transcriptome data increases, the non-parametric and automatic nature of our approach will allow for a corresponding increase in resolution of the analysis resulting in a more detailed description of oscillatory gene expression systems.

## Competing interests

The authors declare that they have no competing interests.

## Author’s contributions

AB and LC did all the computational work. MR, NP and EM conceived the study. All authors participated in writing the paper. All authors have read the paper and approve it.

## Acknowledgements

We thank the Oscope authors for sharing their processed microarray Whitfield data.

## Funding

MR and AB were supported by MRC award MR/M008908/1. LC was supported by a EU Marie Curie fellowship. NP is funded through a Wellcome Trust Senior Research Fellowship (106185/Z/14/Z). EM is funded through a Sir Henry Wellcome Fellowship (201380/Z/16/Z).

## Availability of data and materials

All data used in the paper are freely available; see Leng *et al.* [10] for details on download instructions. The OscoNet code is publicly available at https://github.com/alexisboukouvalas/OscoNet.

## Additional Files

A supplementary report is available (supplementary.pdf).

## Quality control

In any single-cell pipeline, it is necessary to perform some rudimentary quality control to ensure downstream analysis is not affected by low-quality measurements. In the Oscope pipeline [10], a threshold on the mean level and a log-linear cut-off on the variance is applied.

We prefer using a simpler approach based on variance based filtering with a high variance cut-off of 0.9 [5]. This ensures a simple and consistent quality control mechanism that has been shown to remove low quality genes [5].

## Roughness measure

Roughness for a particular gene was defined in [15] as

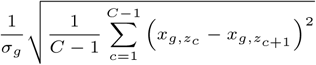

where *σ_g_* the standard deviation of gene expression and *x_g,zc_* is the gene expression for gene *g* at pseudo-time order *z*_*c*_.

This metric measures the smoothness of the gene expression profile by looking at the differences of consecutive measurements. Smaller values indicate a smoother response.

## Footnotes

[1] Intuitively, any monotonic transformation of time will not alter the value of the objective function.

[2] These are available in the *R* package *igraph*.

